# An agent-based modelling framework for tumour growth incorporating mechanical and evolutionary aspects of cell dynamics

**DOI:** 10.1101/2024.05.30.596685

**Authors:** Cicely K Macnamara, I. Ramis-Conde, Tommaso Lorenzi, Alfonso Caiazzo

## Abstract

We develop an agent-based modelling framework for tumour growth that incorporates both mechanical and evolutionary aspects of the spatio-temporal dynamics of cancer cells. In this framework, cells are regarded as viscoelastic spheres that interact with other neighbouring cells through mechanical forces. The phenotypic state of each cell is described by the level of expression of an hypoxia-inducible factor that regulates the cellular response to available oxygen. The rules that govern proliferation and death of cells in different phenotypic states are then defined by integrating mechanical constraints and evolutionary principles. Computational simulations of the model are carried out under a variety of scenarios corresponding to different intra-tumoural distributions of oxygen. The results obtained, which indicate excellent agreement between simulation outputs and the results of formal analysis of phenotypic selection, recapitulate the emergence of stable phenotypic heterogeneity among cancer cells driven by inhomogeneities in the intra-tumoural distribution of oxygen. This article is intended to present a proof of concept for the ideas underlying the proposed modelling framework, with the aim to apply the related modelling methods to elucidate specific aspects of cancer progression in the future.

## 1. Introduction

The term cancer is used to mean any disease in which a population of cells grows to an abnormal degree with the potential to invade the surrounding tissue or spread to other parts of the body. Cancerous cells have many features, *Hallmarks*, which distinguish them from normal healthy cells, for example, sustained proliferation, evasion of apoptosis, and genome instability [16, 17].

Hypoxia-inducible factors (HIFs) e.g. HIF-1*α* [20] are transcription factors that are activated in response to low oxygen levels, i.e. hypoxic conditions. The study of cancerous masses under hypoxic conditions and the role played by HIFs is of particular importance to our understanding of cancer progression. The unrestricted proliferation of cancer cells quickly depletes the available oxygen, particularly in the avascular stage of tumour growth. One key mechanism in response to this is angiogenesis leading to the more malignant vascular stage of tumour growth [17]. Certain cancer treatments (e.g. angiogenesis inhibiting chemotherapies such as thalidomide and bevacizumab) have previously been designed to deprive cancer cells of oxygen with the goal being to inhibit their growth, as hypothesised by Folkman [10]. Unfortunately, on the contrary, drug-induced hypoxia had the opposite effect, instead promoting tumour growth and spread by activating genes involved in angiogenesis, glycolysis, and invasion [39]. Hypoxia-inducible pathways driven by HIFs are responsible for providing cancer cells with an adaptive advantage to hypoxic conditions [29]. Indeed, hypoxic conditions and the subsequent activation of HIFs promote cancer development. Therefore, HIFs are now a target for cancer therapy with research turning to the investigation of the potential of HIF inhibitors [19].

The interplay between variability in cell expression of HIFs, which arises from a variety of genetic and epigenetic mechanisms, and inhomogeneity in oxygen distribution throughout the tumour has been demonstrated to play a key role in the emergence of intra-tumour phenotypic heterogeneity. In more detail, cells that display higher levels of HIFs are found in poorly-oxygenated parts of the tumour (i.e. the interior of the tumour in avascular tumours and regions far from blood vessels in vascularised tumours), while cells with lower levels of HIFs are primarily detected in well-oxygenated parts of the tumour (i.e. the tumour border in avascular tumours and the regions in the vicinity of blood vessels in vascularised tumours) [15, 20, 30, 33, 36, 38, 42]. Such a form of spatial segregation between cells with different levels of HIFs is rooted in the different phenotypic characteristics that are expressed by these cells. In fact, broadly speaking, cells with higher levels of HIFs express a more hypoxic phenotype being better adapted to the environmental conditions of poorly-oxygenated regions of the tumour, whereas cells with lower levels of HIFs express a more normoxic phenotype and, therefore, are better adapted to the environmental conditions of well-oxygenated tumour regions [1, 12, 14, 21, 26, 27].

Mathematical models of tumour growth may be divided roughly into continuum models formulated as differential equations and agent-based models. Continuum models treat cells as a continuous medium and so provide an aggregate description of cell dynamics at the population level. These models are, typically speaking, more computationally efficient and analytically tractable, but they cannot capture the explicit behaviour of individual cells. Agent-based models, on the other hand, treat cells as individual entities and thus can capture detailed cell behaviours more accurately, which makes it possible to describe cell dynamics at the single-cell level, but this frequently comes at a high computational cost and limited analytical tractability. Both continuum models [8, 40] and agent-based models [31, 35] of tumour growth, development, and treatment have been used to study the involvement of HIFs. Other mathematical models have focused on the intra-tumoural role of HIFs within specific Hypoxia-inducible pathways [3, 9] or within entire signalling networks [11, 28]. Continuum models of phenotypic heterogeniety within tumours not specifically related to HIFs have also been considered [6, 5]. Specifically for this paper we are interested in the emergence of phenotypic heterogeneity in HIF expression among individual cells mediated by changes in local oxygen concentration within solid tumours. Such phenotypic heterogeniety was investigated in [40] by means of a continuum model that neglects mechanical aspects of cell dynamics. In this article we aim to investigate it by means of an agent-based model that also captures mechanical interactions of cells.

To this end, we extend the modelling approach for tumour growth that we presented in [4, 25] by developing an agent-based modelling framework that incorporates both mechanical and evolutionary aspects of the spatio-temporal dynamics of cancer cells which underlies tumour growth. Cells are regarded as viscoelastic spheres that interact with other neighbouring cells through mechanical forces, as per [4, 25]. Building on the continuum modelling approach developed in [2, 23, 40], we extend our modelling approach by implementing the dynamics of the phenotypic state of each cell. We choose to describe the phenotypic state by the level of expression of an HIF, whose value can change in time due to random fluctuations. The rules that govern proliferation and death of cells in different phenotypic states are then defined, depending on the availability of space and oxygen, by integrating mechanical constraints and evolutionary principles. We carry out computational simulations of the model, investigating various scenarios corresponding to different intra-tumoural distributions of oxygen. The results obtained show the emergence of stable phenotypic heterogeneity among cancer cells driven by inhomogeneities in the intra-tumoural distribution of oxygen. They indicate excellent agreement between our simulation outputs, the results of [40] and formal analysis of phenotypic selection.

The remainder of the article is organised as follows. In Section 2, we describe our agent-based modelling framework. In Section 3, we provide the main results of computational simulations of the model. In Section 4, we discuss these results and propose some future research directions.

## 2. Modelling Framework

Our modelling framework for tumour growth describes each cancer cell as a viscoelastic sphere that interacts with other neighbouring cells through mechanical forces, as outlined in Section 2.1. The phenotypic state of each cell is allowed to change over time due to random fluctuations, as detailed in Section 2.2. The rules that govern mitosis and death of cells in different phenotypic states are then defined by integrating mechanical constraints and evolutionary principles, as described in Section 2.3.

### 2.1 Mechanical interactions between cells

The position of the centre of cell *i* at time *t* is modelled by the vector function **x**_*i*_(*t*), taking values in the tissue domain where cells are contained. Cell-cell mechanical interactions are then taken into account by letting the evolution of **x**_*i*_(*t*) over a time step Δ*t* be governed by the following discrete equation of motion [25]

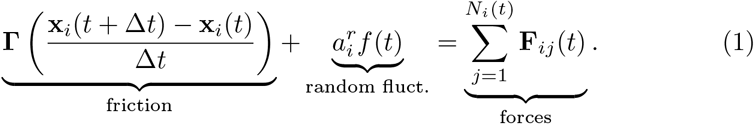

Here, **Γ** is a tensor that models the physical structure of the environment surrounding the cells and can be thought of as tissue friction, for simplicity assumed to be isotropic – i.e. **Γ** = *γ***1**, where *γ* is a positive friction constant and **1** is the identity matrix. The term 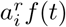 models active random forces exerted by cell *i* as a process of exploration of the nearby space, where *f* is a normal distribution with zero mean and unit variance, and 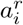 is the amplitude of the random forces [13]. Moreover, *N*_*i*_(*t*) denotes the total number of cells in contact with cell *i* and **F**_*ij*_(*t*) is the force exerted on cell *i* by a neighbouring cell *j*. We let the cell-cell interaction force for cell *i* in contact with cell *j* at time *t* be directed along the vector joining the centres of the two cells, **d**_*ij*_(*t*), and consist of a combination of repulsive and attractive forces. In particular, denoting with *R*_*i*_(*t*) and *R*_*j*_(*t*) the radii of cells *i* and *j* at time *t*, with *E*_*i*_ and *E*_*j*_ the cells’ respective Young’s moduli, and with *ν*_*i*_ and *ν*_*j*_ their Poisson ratios, we use the following definition

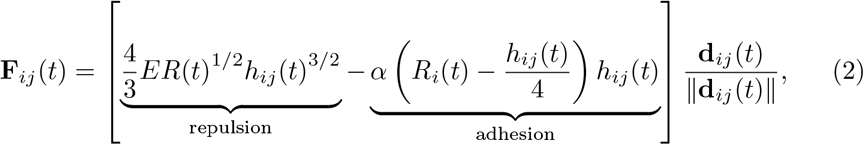

where ∥ · ∥ stands for the Euclidean norm. Here, *R*(*t*) is the effective radius and *E* is the effective Young’s Modulus, which are given by

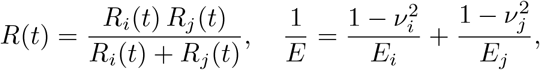

*h*_*ij*_(*t*) is the length of “overlap” (deformation) between cells *i* and *j* at time *t*, which is given by

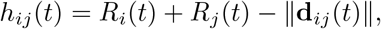

and *α* is the adhesion coefficient. Note that in Equation (2) we assume, without loss of generality, that positive forces for cell *i* are directed outwards. The repulsive force is derived from the Hertz model [18] assuming sufficiently small deformations of the spherical cell, while the adhesion force is assumed to be proportional to the contact surface between two cells and it is computed as per [32]. For more details of the mechanical framework we refer the reader to [4, 25].

### 2.2. Phenotypic changes

The phenotype of cell *i* at time *t* is modelled by a scalar function, *σ*_*i*_(*t*), taking values in a uniform discretisation, of step Δ*σ*, of the interval [0, 1]. We relate the value of *σ*_*i*_(*t*) to the normalised level of a cell’s expression of an HIF − specifically, larger values of *σ*_*i*_ correspond to lower expression of an HIF. We consider phenotypic changes caused by random fluctuations in HIF expression, thus, we allow every cell to update its phenotype according to a random walk. Specifically, over a time step Δ*t*, cell *i* is permitted to acquire either of the phenotypes *σ*_*i*_(*t*) ± Δ*σ* based on a uniform probability distribution that derives the rate of phenotypic change, *β*. Attempted phenotypic changes that imply acquiring a phenotype outside the interval [0, 1] are aborted.

### 2.3. Mitosis and cell death

Mitosis is modelled as the volume preserving splitting of one parent cell into two equal-sized progeny cells, which take on the phenotype of the parent cell, over a time step Δ*t*. It is governed by necessary biological and mechanical conditions along with evolutionary conditions involving the fitness of the cell’s phenotype. Firstly, cells are only allowed to undergo mitosis after reaching a mature state, considered to be when the cell has grown to at least 99% of a prescribed maximum radius, *R*_*M*_. The cell cycle is modelled by assuming that each cell increases in size at a given growth rate, *θ*, until it reaches its maximum size. Mitosis, however, is not possible if the cell experiences an excessive compression force due to neighbouring cells (contact inhibition). To take this into account, mitosis is only allowed as long as (i) the repulsive force of the modified Hertz model (see Section 2.1) is below a given threshold, *H*^*T*^, and (ii) the total number of contact neighbours of the cell is below a given threshold, *N* ^*T*^.

Moreover, we incorporate into the model the effect of environmental selection on HIF expression by postulating that if the aforementioned necessary biological and mechanical conditions are met then the rate of mitosis is determined by the cell’s fitness, which depends on the cell’s phenotype. In more detail, denoting by ℛ_*i*_(*t*) the fitness of cell *i* with phenotype *σ*_*i*_(*t*) at time *t*, if ℛ_*i*_(*t*) > 0 then over a time step Δ*t* we let cell *i* undergo mitosis with probability 0 < Δ*t* ℛ_*i*_(*t*) ≤ 1. Similarly, we let cells die at a rate determined by the cell’s fitness. Specifically, if R _*i*_(*t*) < 0 then over a time step Δ*t* we let cell *i* die with probability 0 < Δ*t* |R_*i*_(*t*) | ≤ 1. Dead cells are instantaneously removed from the system.

Building upon [2, 23, 40], we use the following definition of the fitness of cell *i* at time *t*

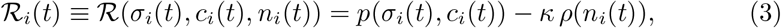

which depends on the phenotype of the cell, *σ*_*i*_(*t*), the concentration of oxygen local to the cell, *c*_*i*_(*t*), and the number of cells in a particular spatial location close to the cell, *n*_*i*_(*t*). Our model is completely off-lattice, however, for computational efficiency we split the spatial domain into cube (square, in 2D) units of size 10*μm*^3^ (10*μm*^2^, in 2D) to restrict neighbour searching to the cube of the cell and surrounding cubes. For the purposes of this work, we consider *n*_*i*_(*t*) to be the number of cells in the same cube unit as cell *i*.

The function *ρ*(*n*_*i*_) in Equation (3) captures the rate of cell death induced by competition for space (i.e. saturated growth), and it is thus a non-negative and monotonically increasing function, i.e.

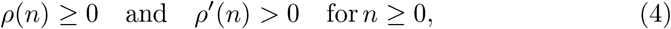

while the parameter *κ* > 0 measures the impact of saturated growth on the cell fitness.

The function *p*(*σ*_*i*_, *c*_*i*_) in Equation (3) captures the intrinsic division rate of cell *i* with phenotype *σ*_*i*_ under the environmental conditions corresponding to the oxygen concentration *c*_*i*_. Following [2, 23, 40], we define

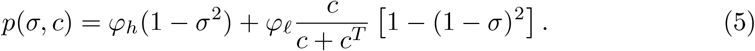

The definition given by Equation (5) relies on the following ideas: (i) cells with phenotype *σ* = 1 (i.e. cells with the lowest normalised level of HIF expression) have a fully oxidative metabolism and produce the energy required for mitosis through aerobic respiration only; (ii) cells with phenotype *σ* = 0 (i.e. cells with the highest normalised level of HIF expression) have a fully glycolytic metabolism and produce energy through anaerobic glycolysis only; (iii) cells with other phenotypes *σ* ∈ (0, 1) (i.e. cells with intermediate normalised levels of HIF expression) produce energy via both aerobic respiration and anaerobic glycolysis, and smaller values of *σ* (i.e. higher normalised levels of HIF expression) correlate with a less oxidative and more glycolytic metabolism. Here *c*^*T*^ > 0 is the half-saturation constant of oxygen and *φ*_*h*_ > 0 and *φ*_*ℓ*_ > 0 are the intrinsic division rates associated with the highest and lowest normalised level of HIF expression, respectively. They should not be taken to be the actual rates of proliferation of cells since, as discussed earlier in this Section, the proliferation of cells is also determined by other biological and mechanical conditions. Indeed the resulting cell cycle rate is much lower. As noted in [2], Equation (5) leads to a fitness function that is close to the approximate fitness landscapes which can be inferred from experimental data through regression techniques.

As per the formal analysis of phenotypic selection carried out in [40] and recapitulated in Appendix A, under the definition of the fitness function given by Equations (3) and (5), if the oxygen concentration is fixed at some constant value 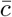 (i.e. 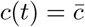 for all *t* ≥ 0) in a certain region of the tumour tissue, then the phenotype that can be expected to be ultimately selected in that region (i.e. the expected phenotype) is given by

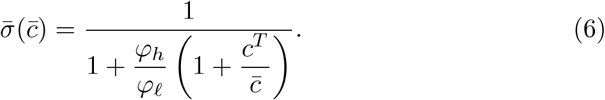

Note that, in accordance with biological evidence [15, 30, 33, 38, 42], the expression of the expected phenotype given by Equation (6) is such that: 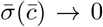 as 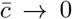, indicating that cells with the highest normalised level of HIF expression can be expected to be selected in hypoxic regions of the tumour; 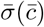 attains larger values as 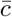 increases, indicating that cells with lower levels of HIF expression can be expected to be selected in more oxygenated tumour regions.

Finally, we note, that cells which are created or move out of the domain along the domain boundary are removed instantly. This is an acceptable simplification since we can make the domain arbitrarily large and we are not interested in behaviours which occur at the edge of the domain.

## 3. Main Results of Computational Simulations

In this section, we present the main results of computational simulations of the agent-based model described in the previous section, under a variety of scenarios corresponding to different intra-tumoural distributions of oxygen.

### 3.1. Simulation setup

Simulations were carried out using the parameter values provided in Tables 1 and 2, which were chosen in line with those in [4, 25, 40] and references therein^1^. Specific differences between the values in [40] and Table 2 relate to a computational choice here to allow phenotypic changes to occur over the same mechanics timestep used throughout the code. A future improvement could rely on different timesteps for the mechanics and evolutionary aspects.

**Table 1:** Parameter values used to simulate mechanical interactions between cells. These values were chosen in line with those in [4, 25].

| Parameter | Description | Value |
| --- | --- | --- |
| $\Delta t$ | timestep | 1 min |
| $R_i$ | cell radius | $3.97 - 5 \mu\text{m}$ |
| $E_i$ | Young's modulus | $1 \times 10^{-3} \mu\text{N}/\mu\text{m}^2$ |
| $\nu_i$ | Poisson ratio | 0.5 |
| $\alpha$ | adhesion coefficient | $3.72 \times 10^{-4} \mu\text{N}/\mu\text{m}^2$ |
| $\gamma$ | cell-medium friction constant | $1 \times 10^{-2} \mu\text{N}\mu\text{m}/\text{min}$ |
| $a_i^r$ | amplitude of random forces | $4 \times 10^{-3} \mu\text{N}$ |

**Table 2:** Parameter values used to simulate phenotypic changes, mitosis, and cell death. These values were chosen in line with those in [4, 25, 40].

|  | Description | Value |
| --- | --- | --- |
| $\Delta\sigma$ | phenotype interval stepsize | $5 \times 10^{-3}$ |
| $\theta$ | growth rate of cell size | $0.1 \mu\text{m}/\text{min}$ |
| $R_M$ | cell maximum radius | $5 \mu\text{m}$ |
| $H^T$ | threshold repulsive force above which mitosis is inhibited | $5 \times 10^{-4} \mu\text{N}$ |
| $N^T$ | number of contact neighbours above which mitosis is inhibited | 8 in 2D, 16 in 3D |
| $\beta$ | rate of phenotypic change | $1 \times 10^{-5} / \text{min}$ |
| $\varphi_h$ | intrinsic division rate of cells with the highest HIF expression | $1 \times 10^{-2} / \text{min}$ |
| $\varphi_\ell$ | intrinsic division rate of cells with the lowest HIF expression | $1 \times 10^{-1} / \text{min}$ |
| $c^T$ | half-saturation constant of O2 | 50 mmHg |
| $\kappa$ | impact of saturated growth on cell fitness | $1.1 \times 10^{-4} / \text{min}$ |

Moreover, we used the following definition of the function *ρ*(*n*) in Equation (3)

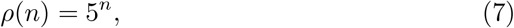

which is informed by the results of preliminary simulations that we carried out considering different definitions of the function *ρ*(*n*). In particular, we investigated the case where *ρ*(*n*) = *n* and the case where *ρ*(*n*) = *a*^*n*^ with *a* > 1. Among the definitions of the function *ρ*(*n*) that we considered, the one given by Equation (7) leads, under the parameter values given in Tables 1 and 2, to a fitness function ℛ(*σ, c, n*) defined via Equation (3) that can take both positive and negative values – i.e. a fitness function that allows both mitosis and cell death to occur depending on the cell phenotype *σ* – for biologically relevant values of the oxygen concentration *c*, under the constraints on the cell number *n* that are imposed by the biological and mechanical conditions described in Section 2.1. Cell death is an important feature as it allows space for mitosis throughout the simulated time and it is this continued mitosis, through the function given in Equation 5, which drives phenotypic selection. We note that in the first three simulations detailed here, during the initial phase a single cell starts a cascade of proliferation growing a population of cells which eventually fills the entire tissue domain. After this tissue growth phase is where we can appreciate changes in the phenotypic composition of the tumour due to the changeover in the cell population.

### 3.2. Simulations under fixed homogeneous oxygen distribution

We begin by considering a 2D tissue domain, measuring 800*μ*m × 800*μ*m, exposed to a fixed homogeneous oxygen distribution (i.e. the oxygen concentration is kept fixed to some value 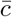 throughout the whole tissue domain). We place a single cell, of phenotype 0.5, at the centre of the domain at the start and carry out simulations for different values of 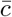 – namely 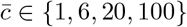 mmHg − using the parameters given in Tables 1 and 2. As per Equation (6), the resulting expected phenotypes for the four different oxygen concentrations are 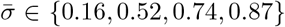 (2dp), respectively (recall, *c*^*T*^ = 50mmHg). The results obtained are shown in Figure 1 in panels 1a, 1b, 1c and 1d respectively. The expected phenotype values are indicated by the horizontal red dashed lines in each plot of Figure 1. The black lines in Figure 1 indicate the average phenotype of all cancer cells in the domain at a given timestep, while the grey shaded regions indicate the corresponding standard deviation. For each case we observe that the average phenotype tends to the expected phenotype after sufficiently long time. Moreover, as indicated by the grey shaded regions, there is small variability between cells’ phenotypes. Plots showing the spatial distribution of cells across the 2D domain at time intervals are given in Appendix B confirming the small variability, and thus, homogeneity, between cell phenotypes across the domain.

**Figure 1.**
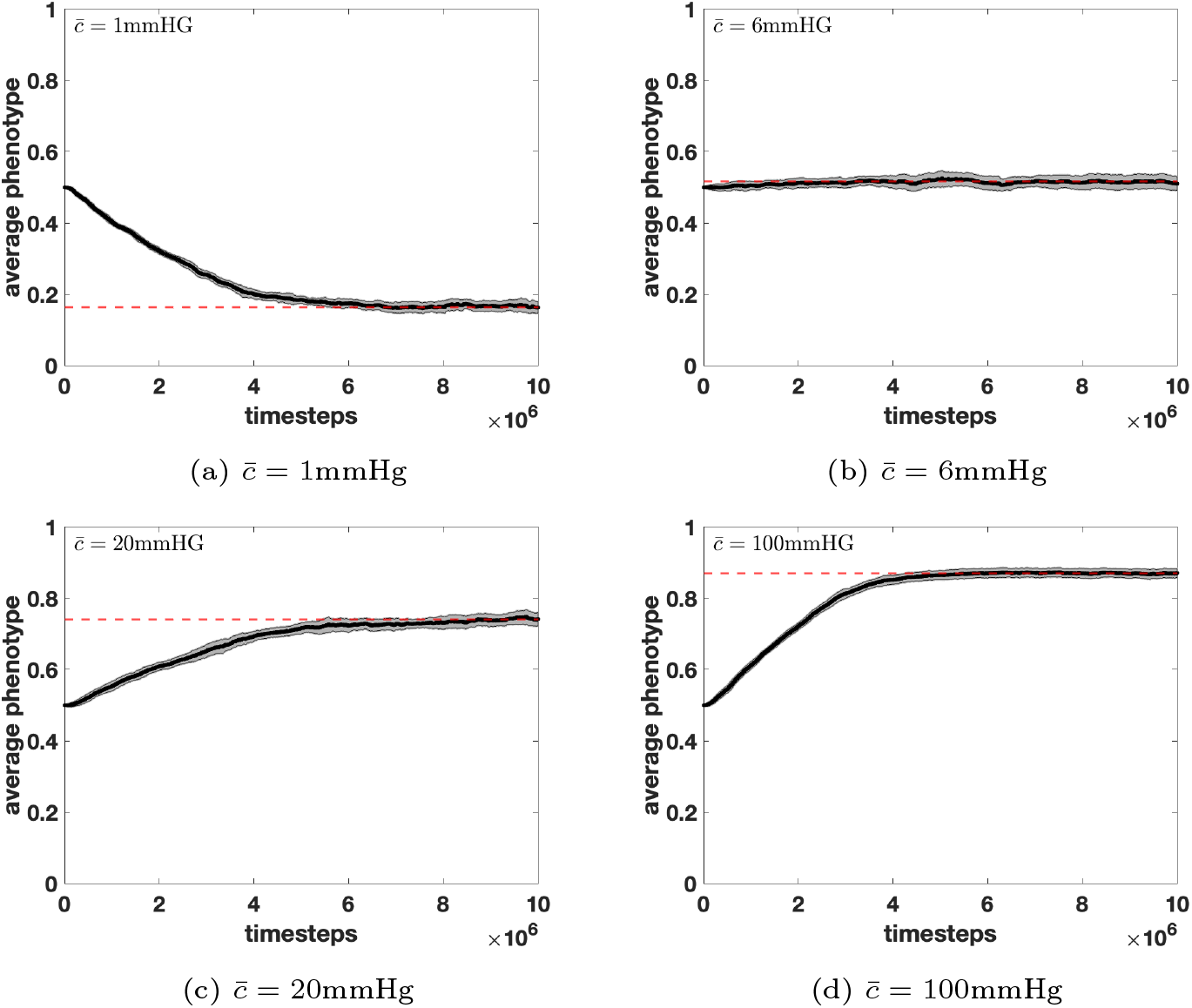
Simulations under fixed homogeneous oxygen distribution. Plots showing the mean phenotype of cancer cells (black line) along with the corresponding standard deviation (grey shaded region) over time for a 2D tissue domain where the oxygen concentration is kept fixed to some value 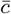 throughout the whole domain. Four different values of 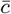 are considered, as specified in the top left corner of each panel – namely 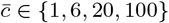 mmHg, in panels (a), (b), (c) and (d), respectively. The red dashed lines show the expected phenotype as given by Equation (6). These plots correspond to the parameter values given in Tables 1 and 2.

### 3.3. Simulations under fixed heterogeneous oxygen distribution

We then consider a 3D tissue domain, measuring 200*μ*m× 200*μ*m ×200*μ*m, exposed to a fixed heterogeneous oxygen distribution (i.e. the oxygen concentration is kept fixed to different values in different tumour regions). The purpose of these experiments is to show that cells, allowed to change their phenotype completely randomly, nonetheless exhibit spatial segregation, with cells eventually matching their phenotype to the local conditions. Once again a single cell with a phenotype of 0.5 is placed in the centre of the domain at the start of the simulation.

We first investigate a scenario where the oxygen concentration is fixed to two different values in two equal halves of the domain. In particular, we choose the left-hand side of the domain to have a fixed oxygen level of 1mmHg and the right-hand side of the domain to have a fixed oxygen level of 100mmHg. The results of simulations are shown in Figure 2. In panel 2d we colour the cells according to their local concentration of oxygen showing the imposed spatial segregation of oxygen. In panel 2f we show the phenotypes expressed by the cells at the end of the simulation. We observe that for both halves of the domain the majority of cells express the expected phenotypes given by Equation (6) – i.e. the phenotype 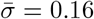 on the left-hand side of the domain, where the oxygen concentration is kept constant to 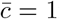 mmHg, and the phenotype 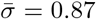 on the right-hand side of the domain, where the oxygen concentration is kept constant to 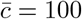 mmHg. At the boundary between the two halves of the domain, we find cells of either phenotype. We arbitrarily split the spatial domain into three equal sections, so as to further consider the boundary of the two halves of the domain, as indicated in panel 2e. We plot the average phenotypes of cancer cells in these regions over time, as shown in panels 2a, 2b, and 2c. The expected phenotypes are achieved after a sufficient amount of time in the left (panel 2a) and right (panel 2c) thirds of the domain. In the middle third of the domain (panel 2b) there is a mixture of cells of both phenotypes, specifically along the boundary plane of the two halves of the domain, leading to an average phenotype that lies between the two expected values and a much greater standard deviation. The random motility of cells and mechanical interactions cause cells of either of the two expected phenotypes to cross into the other half of the domain continually during the simulations. Since mechanical effects happen far quicker than phenotypic selection there are always cells along the boundary plane which express the phenotype expected for the other half of the domain. Note, there are slightly more cells with phenotype closer to 1 (i.e. a lower expression of an HIF) in this region since the division rate of cells with phenotype closer to 1 is higher. This highlights an aspect which is only captured by nature of this being an agent-based model.

**Figure 2.**
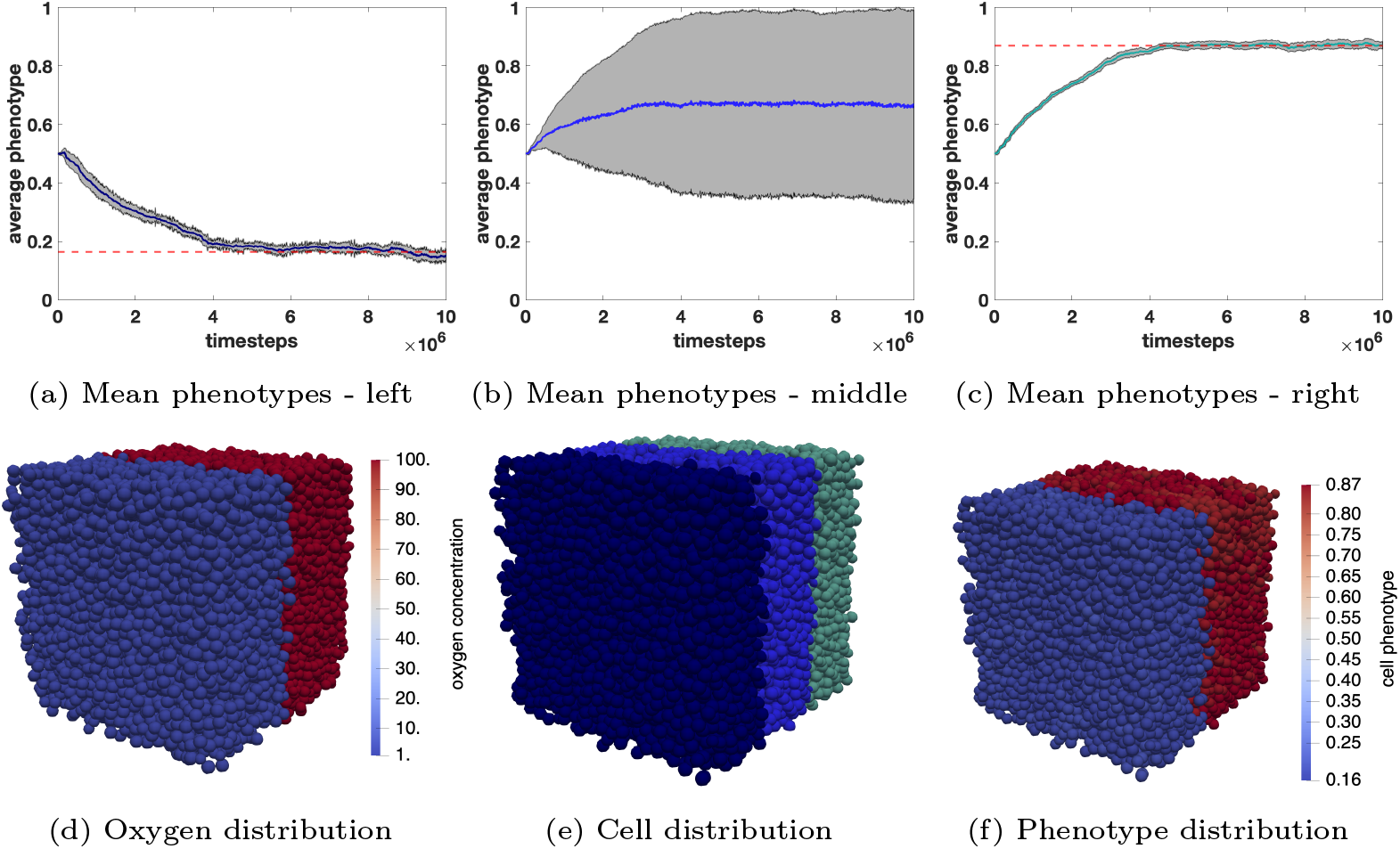
Simulations under fixed heterogeneous oxygen distribution. Plots showing simulation results for a 3D tissue domain where the oxygen concentration is fixed to two different values in two equal halves of the domain – namely the left-hand side of the domain has a fixed oxygen level of 1mmHg and the right-hand side of the domain has a fixed oxygen level of 100mmHg. The plots in panels (a), (b) and (c) show the mean phenotype of cancer cells (dark blue/blue/cyan line, respectively) along with the corresponding standard deviation (grey shaded region) over time for the three thirds of the tumour region highlighted in panel (e). The red dashed lines in panels (a) and (c) show the expected phenotype as given by Equation (6) for the left and right halves of the domain, respectively. The plot in panel (d) shows the spatial distribution of cells at the end of simulations with cells coloured according to their local concentration of oxygen, while the plot in panel (f) shows the spatial distribution of cells at the end of simulations with cells coloured by their phenotype. These plots correspond to the parameter values given in Tables 1 and 2.

We then investigate a scenario where the oxygen concentration is fixed to 100mmHg in a central cylinder (of radius 25*μ*m) of the domain and to 1mmHg everywhere else in the domain. The results obtained are shown in Figure 3 and are analogous to those shown in Figure 2. These results indicate that stable phenotypic heterogeneity of HIF expression is directly driven by inhomogeneities in local oxygen concentration.

**Figure 3.**
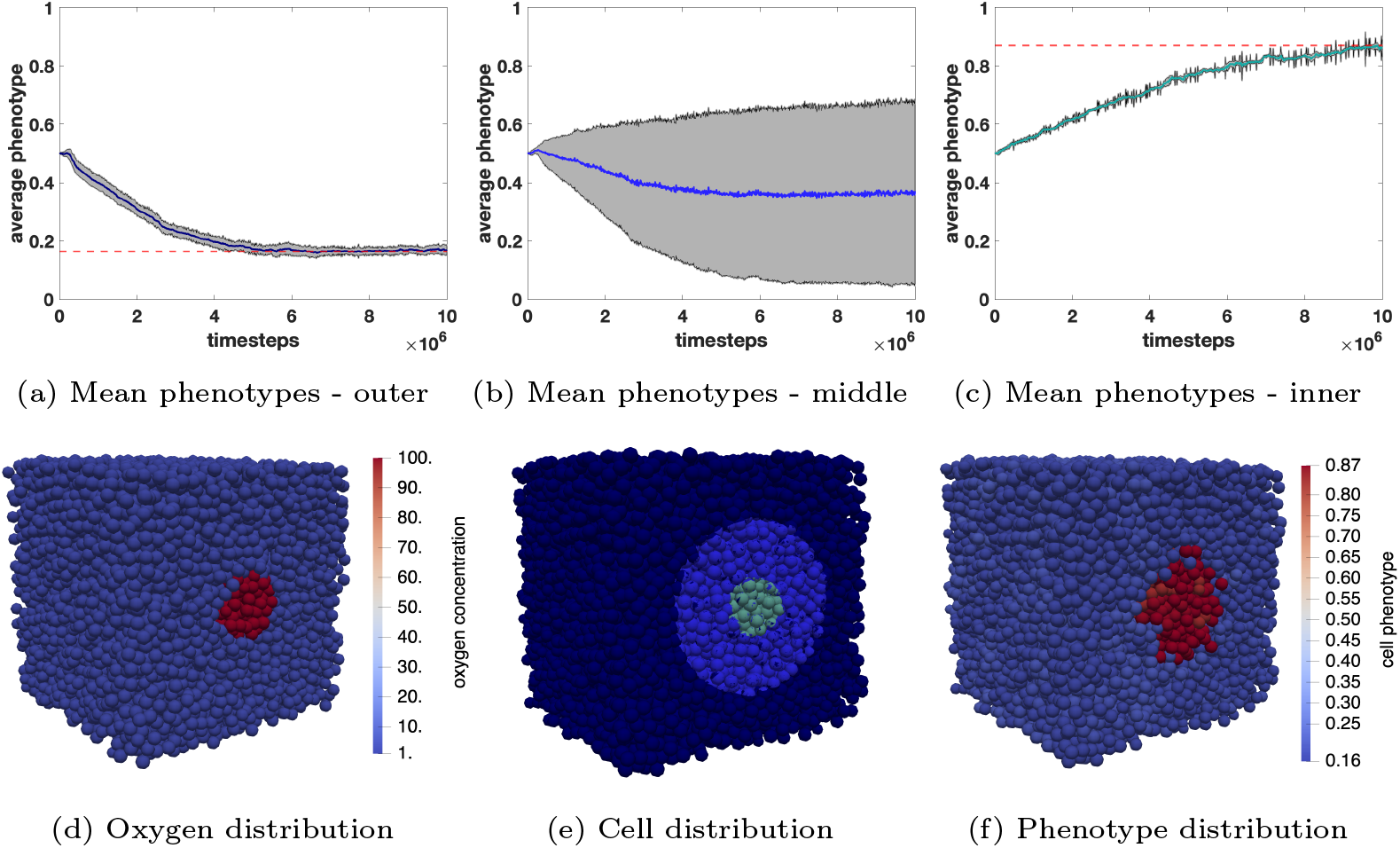
Simulations under fixed heterogeneous oxygen distribution. Plots showing simulation results for a 3D tissue domain where the oxygen concentration is fixed to 100mmHg in a central cylinder of the domain and to 1mmHg everywhere else in the domain. The plots in panels (a), (b) and (c) show the mean phenotype of cancer cells (dark blue/blue/cyan line, respectively) along with the corresponding standard deviation (grey shaded region) over time for the three regions of the tumour domain highlighted in panel (e). The red dashed lines in panels (a) and (c) show the expected phenotype as given by Equation (6) for the exterior and interior of the cylindrical region, respectively. The plot in panel (d) shows the spatial distribution of cells at the end of simulations with cells coloured according to their local concentration of oxygen, while the plot in panel (f) shows the spatial distribution of cells at the end of simulations with cells coloured by their phenotype. These plots correspond to the parameter values given in Tables 1 and 2.

### 3.4. Simulations under dynamically varying heterogeneous oxygen distribution

Finally, we consider a 3D tissue domain of size 400*μ*m × 400*μ*m × 400*μ*m wherein, initially, the bulk of the tissue has a low oxygen concentration (i.e. 1mmHg), but there is a cylindrical region (of radius 50*μ*m) to the bottom right of the domain which has a high oxygen concentration (i.e. 100mmHg). At the start of the simulation we let the cells be distributed uniformly throughout the domain and we assume all cells to have phenotype 0.5. We run the simulation and allow time for the expected phenotypes predicted by Equation (6) to emerge. After such time we introduce a second cylindrical region (again of radius 50*μ*m) of high oxygen concentration (i.e. 100mmHg) to the top left of the domain; we then allow again time for the expected phenotypes to emerge. The obtained results shown as 2D slices in Figure 4 demonstrate that, in analogy with the results shown in Figures 2 and 3, cells with phenotypes closer to 1 are dynamically selected in highly oxygenated regions of the tissue, whereas cells with phenotypes closer to 0 are dynamically selected in poorly oxygenated regions. This indicates that time-variation in the local oxygen concentration leads to dynamical changes in the phenotypic distribution of the cells.

**Figure 4.**
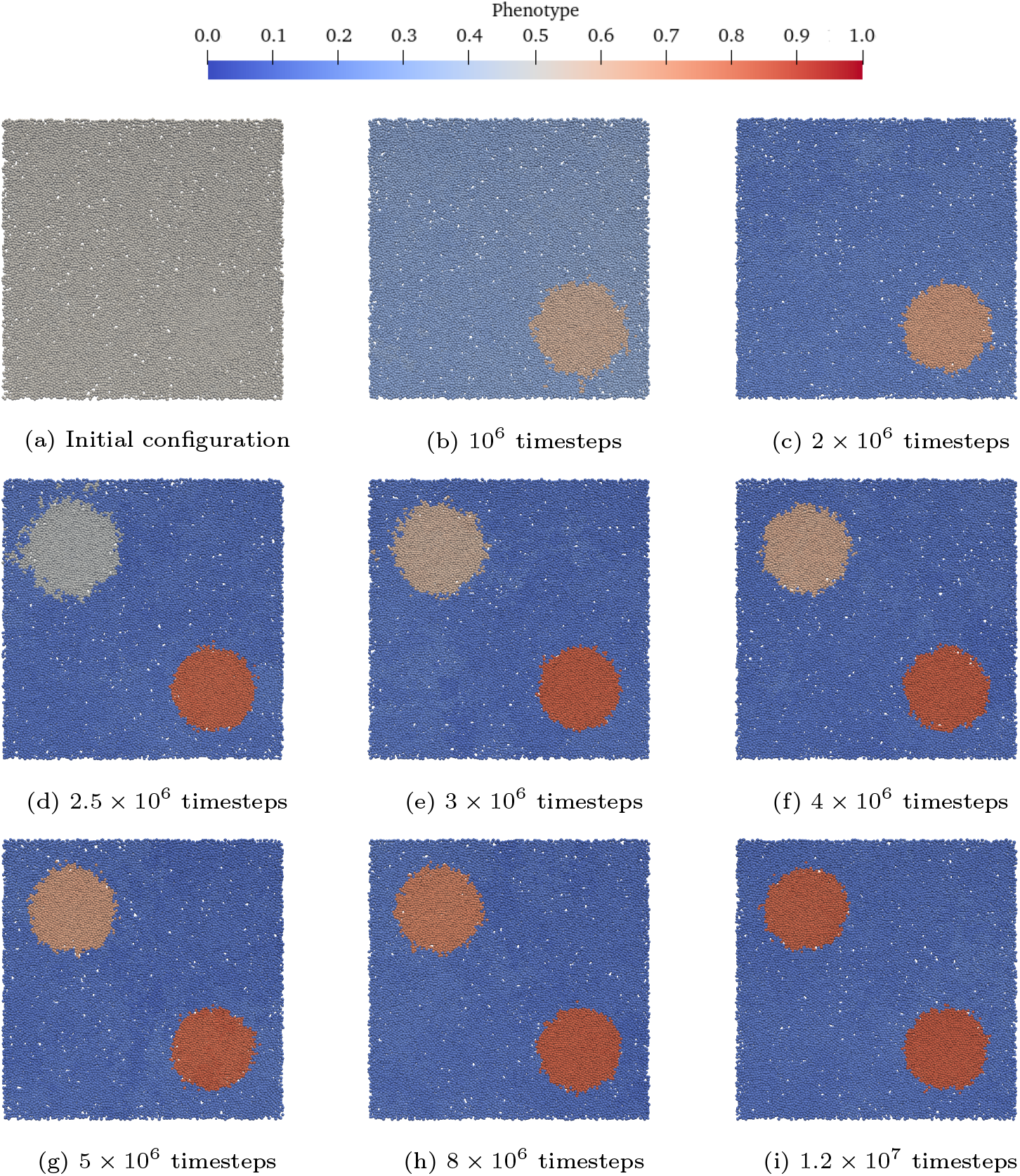
Simulations under dynamically varying heterogeneous oxygen distribution. Plots showing a 2D slice of simulation results for a 3D tissue domain where the oxygen concentration is initially fixed to 100mmHg in a cylindrical region to the bottom right of the domain and to 1mmHg everywhere else in the domain. After 2 ×10^6^ timesteps, a second cylindrical region of oxygen concentration 100mmHg is introduced to the top left of the domain. The different panels display the spatial distribution of cells at different times (specified below each panel), with cells coloured by their phenotype. These plots correspond to the parameter values given in Tables 1 and 2.

## 4. Conclusions and Research Perspectives

We have extended our previous work in [4, 25] by developing an agent-based modelling framework for tumour growth that incorporates both mechanical and evolutionary aspects of cell dynamics. In this framework: (i) cells are regarded as viscoelastic spheres that interact with other neighbouring cells through mechanical forces; (ii) the randomly fluctuating phenotypic state of each cell is described by the level of expression of an HIF; (iii) the rules governing division and death of cells in different phenotypic states are formulated by integrating mechanical constraints [4, 25] and evolutionary principles [2, 23, 40].

In agreement with previous theoretical and empirical work [1, 12, 14, 21, 26, 27], the results of computational simulations of the model indicate that, in the presence of inhomogeneities in the intra-tumoural distribution of oxygen, stable phenotypic heterogeneity among cancer cells can emerge as a result of the effect of environmental selection on HIF expression, leading cells with a lower expression of an HIF to colonise well-oxygenated regions of the tumour tissue, while cells with a higher expression of an HIF outcompete other cells in poorly oxygenated regions. We have also shown that there is an excellent agreement between these simulation outputs and the results of formal analysis of phenotypic selection, which supports the robustness of the obtained computational results.

Although the relationship between cell phenotype, as described by the level of expression of an HIF, and oxygen concentration has been shown in other studies [e.g. 40], the novelty here is the incorporation of these evolutionary aspects into a fully off-lattice agent-based mechanical model for tumour growth and development. As a proof of concept we have shown that this modelling framework can capture the expected biological behaviour in which cells with a low expression of HIF colonize well oxygenated regions of the tumour (and vice versa). The model could be extended to investigate phenotypic variability arising from a range of scenarios.

Building on the modelling approach proposed in [25], the modelling framework presented here could be generalised by introducing an intra-tumour vascular network and incorporating the effect of mechanical interactions between cells and intra-tumour blood vessels. In such a generalised modelling framework, blood vessels would perfuse oxygen into the tumour tissue, which would in turn be taken up by the cells. The dynamics of the oxygen concentration would then be described through a reaction-diffusion equation, which would be coupled with the agent-based model for the dynamics of the cells and would be solved numerically using finite element methods.

Moreover, while we carried out computational simulations under the assumption that cells with different phenotypes share the same mechanical properties (cf. the parameter values in Table 1), phenotypic heterogeneity can drive differences in the mechanical characteristics of the cells, and thus in the values of the parameters implicated in the modelling of cell-cell mechanical interactions. Hence, it would certainly be interesting to explore if incorporating such differences can lead to the emergence of different forms of phenotypic structuring and/or to the formation of different patterns of tumour growth. For example, since it has been reported that cells with a higher level of HIF expression may be more invasive [34, 37, 41], one could assume cells with phenotypes *σ*_*i*_ closer to 0 to be characterised by larger values of the model parameter 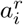 (i.e. the parameter representing the amplitude of active random forces that are exerted by the cells as a process of exploration of the nearby space), so as to make these cells more invasive.

A future improvement of the framework would be to have two working timesteps within the code – one for the mechanical aspects and one for the evolutionary processes. Using the same timestep here, for simplicity, means that our results do not show accurately how long the evolutionary processes take but rather show that given sufficiently long time stable phenotypic structuring will emerge.

Finally, it would be interesting to generalise the limiting procedure employed in [7, 22, 24] expanding the formal analysis in Appendix A in order to formally derive the continuum counterpart of the agent-based modelling framework presented here, which we would expect to consist of a partial integro-differential equation for the local cell population density function (i.e. the local phenotypic distribution of cells). This would make it possible to integrate the computational outputs of the agent-based model with analytical and numerical results on the qualitative and quantitative properties of the solutions of the corresponding continuum model, which would facilitate exploration of the model parameter space, thus permitting a precise identification of the validity domain of the biological predictions made through simulations of the agent-based model.

## Appendix A Formal analysis of phenotypic selection

Following [2, 23, 40], with a little algebra we rewrite the definition of the fitness function given by Equations (3) and (5) as

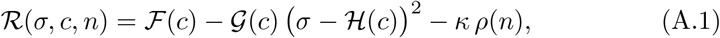

with

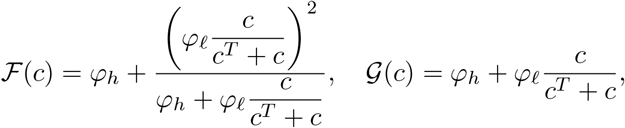

and

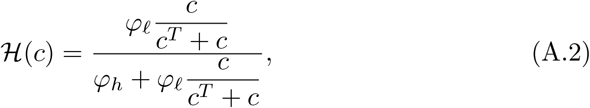

where we have dropped the subscript *i* and the dependence on *t* as the calculations apply independently of the specific cell and time considered. From Equation (A.1) we see that the phenotype with the maximum fitness (i.e. the fittest phenotype) under the environmental conditions corresponding to the oxygen concentration *c* is given by the function ℋ(*c*) defined via Equation (A.2). Hence, if the oxygen concentration is fixed at some constant value 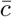 (i.e. 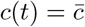 for all *t* ≥0) in a certain region of the tumour tissue, then we expect the phenotype

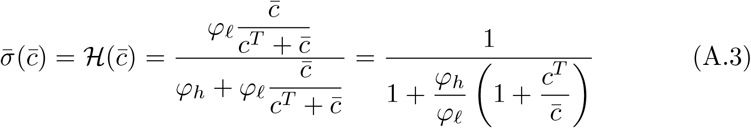

to be ultimately selected in that region. In Figure A.5 we plot the expected phenotype 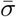 given by Equation (A.3) as a function of the constant oxygen concentration 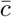 varying in the range 0 − 100mmHg, for three different values of the half-saturation constant of oxygen, *c*^*T*^, under the values of the parameters *φ*_*h*_ and *φ*_*ℓ*_ provided in Table 2.

**Figure A.5:**
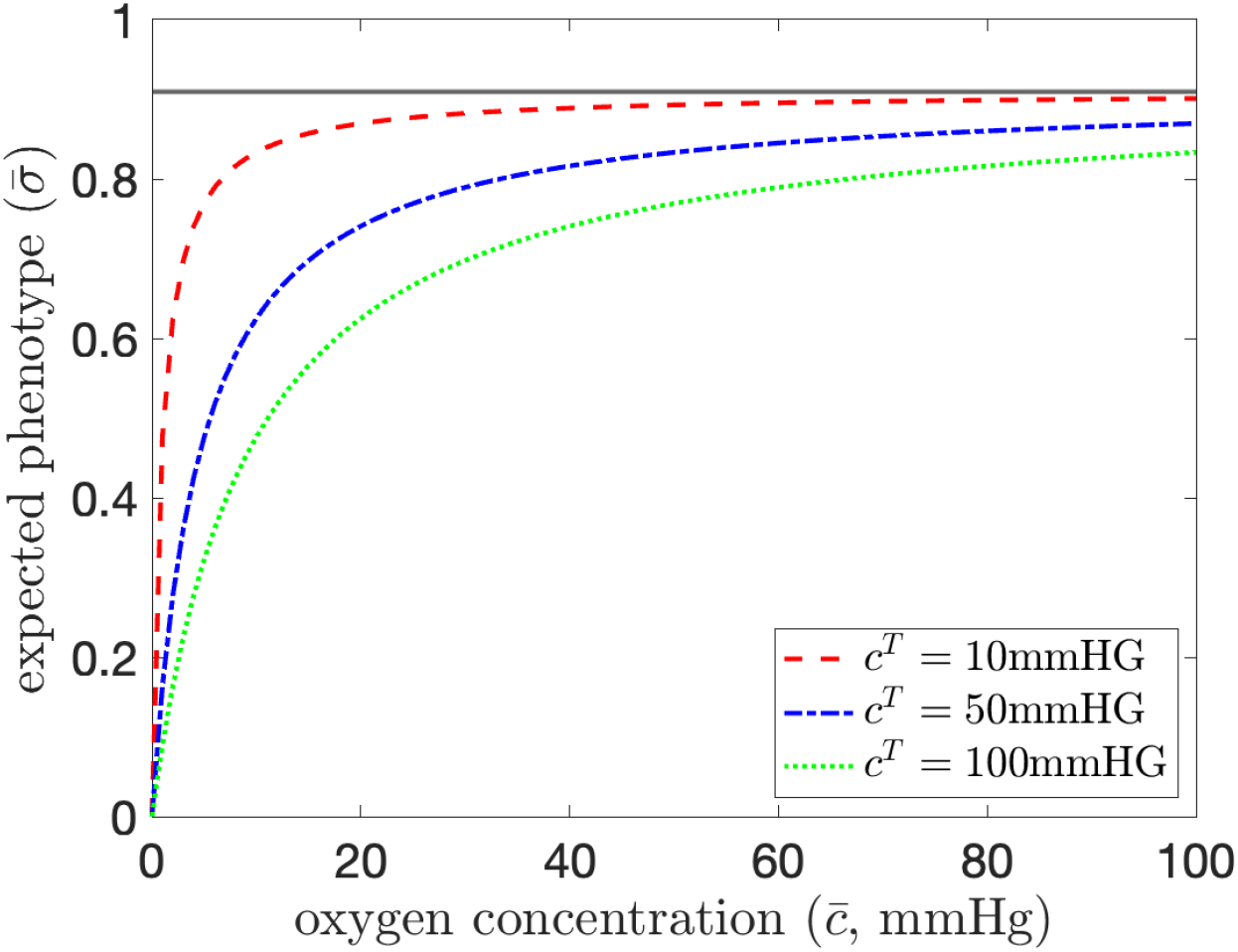
Expected phenotype. Plot of the expected phenotype 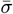, given by Equation (A.3), with respect to the oxygen concentration 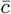 for three different values of the half saturation constant of oxygen, *c*^*T*^, (see legend) under the values of the parameters *ϕ*_*h*_ and *ϕ*_.*ℓ*_ provided in Table 2. The grey horizontal line indicates the corresponding maximum value of the expected phenotype – i.e. the value 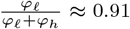– which is attained in the limiting scenario where 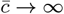.

## Appendix B Cell spatial distribution plots in support of Figure 1

We provide an expansion of the results shown in Figure 1 by providing snapshots in time of the spatial distribution of cells across the whole 2D domain, coloured by phenotype. The snapshots are provided for the cases 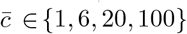 mmHg at 10 000 timesteps which can be thought of as the initial configuration of the tissue as this captures the end of the initial phase of the simulation when the single cell has proliferated into a mass of cells covering the entire domain. We note that it can be observed from Figures B.6b, B.6f, B.6j and B.6n that the density of cells across the tissue domain is affected by the local concentration of oxygen since the proliferation of cells is related to the concentration of oxygen, as per Equation (5). Further snapshots are given after 2×10^6^ timesteps and 4 ×10^6^ timesteps showing the progression of the phenotypes. We observe little variation in the phenotypes across the domain in each case. This confirms that without local variation in oxygen, HIF expression phenotypes are largely homogeneous.

**Figure B.6:**
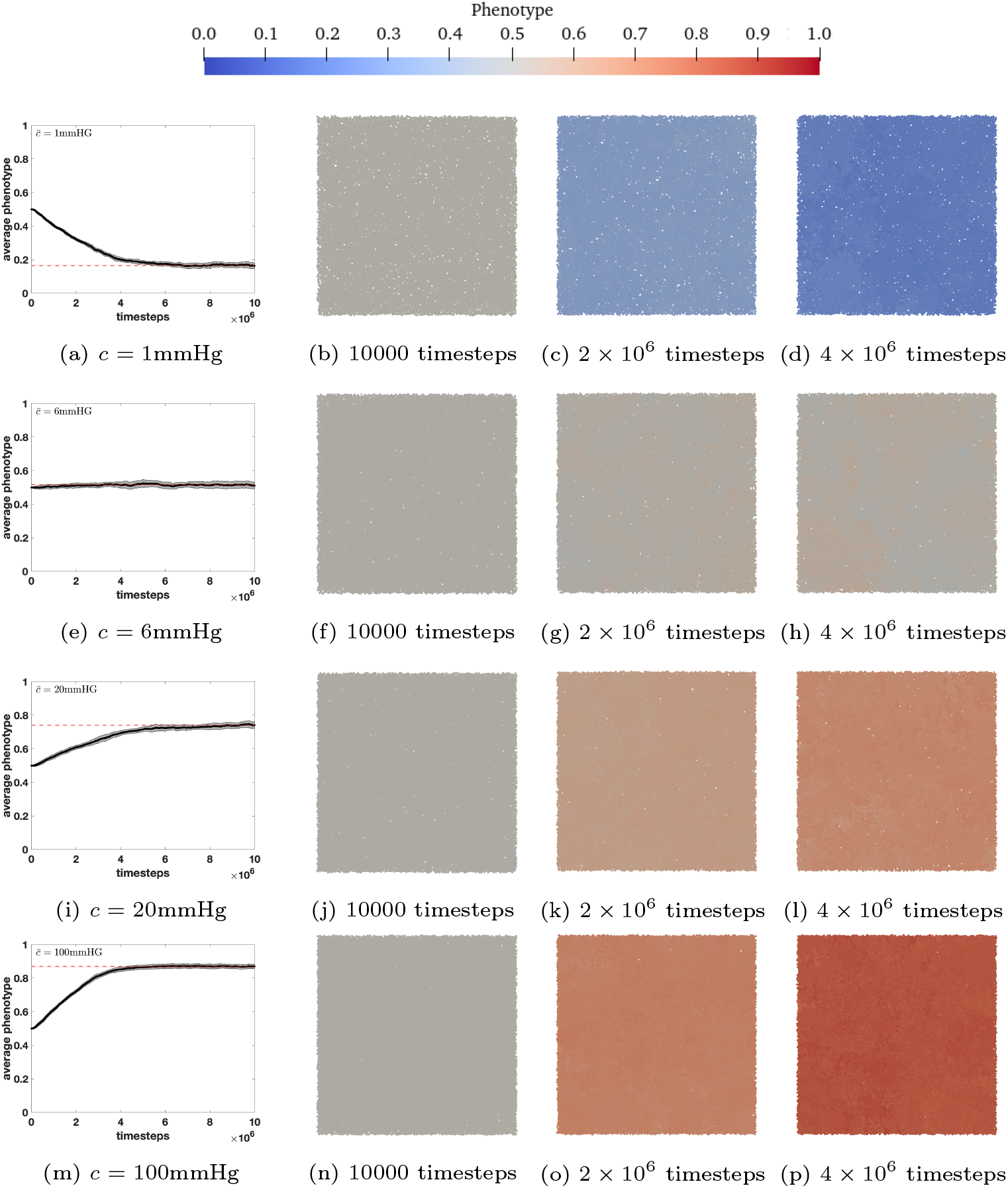
Simulations under fixed homogeneous oxygen distribution. Plots showing the spatial distribution of cells for the 2D simulation results given in Figure 1 and repeated in the left panels (a), (e), (i) and (m) here. Four different values of 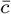 are considered, as specified in the captions for the left panels – namely 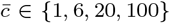 mmHg. We show results after the initial proliferation phase in the panels second from the left panels (b), (f), (j) and (n), after 2 ×10^6^ timesteps, in the third panels (c), (g), (k) and (o) and after 4 × 10^6^ timesteps in the fourth panels (d), (h), (l) and (p). These plots correspond to the parameter values given in Tables 1 and 2.

## Acknowledgements

CKM gratefully acknowledges the support of her Rankin-Sneddon Fellowship at the University of Glasgow. TL gratefully acknowledges support from the Italian Ministry of University and Research (MUR) through the grant PRIN 2020 project (No. 2020JLWP23) “Integrated Mathematical Approaches to Socio-Epidemiological Dynamics” (CUP: E15F21005420006) and the grant PRIN2022-PNRR project (No. P2022Z7ZAJ) “A Unitary Mathematical Framework for Modelling Muscular Dystrophies” (CUP: E53D23018070001) funded by the European Union - Next Generation EU, and from the Istituto Nazionale di Alta Matem-atica (INdAM) and the Gruppo Nazionale per la Fisica Matematica (GNFM).

## Footnotes

1 Note that, since in the modelling framework presented here cells may have different phenotypes, we could have altered the mechanical properties of the cells in line with their level of HIF expression. However, since this is not directly related to the purpose of this study, we decided to save it for future work. Hence, for simplicity, we consider the mechanical properties of each cell to be equivalent (i.e. *E*_*i*_ =1 × 10^−3^*μ*N/*μ*m^2^ and *ν*_*i*_ = 0.5 for all *i*) and that all cells exert random forces of the same amplitude (i.e. 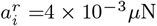 for all *i*).

## Notes

### Competing Interest Statement

The authors have declared no competing interest.

### Summary of Updates

Some slight changes to the text to make things clearer and sent to a different journal.

